# On the Reliability of Chronically Implanted Thin-Film Electrodes in Human Arm Nerves for Neuroprosthetic Applications

**DOI:** 10.1101/653964

**Authors:** P. Čvančara, G. Valle, M. Müller, T. Guiho, A. Hiairrassary, F. Petrini, S. Raspopovic, I. Strauss, G. Granata, E. Fernandez, P. M. Rossini, M. Barbaro, K. Yoshida, W. Jensen, J.-L. Divoux, D. Guiraud, S. Micera, T. Stieglitz

## Abstract

Direct stimulation of peripheral nerves can successfully provide sensory feedback to amputees while using hand prostheses. Recent clinical studies have addressed this important limitation of current prostheses solutions using different implantable electrode concepts. Longevity of the electrodes is key to success. We have improved the long-term stability of the polyimide-based transverse intrafascicular multichannel electrode (TIME) that showed promising performance in clinical trials by integration of silicon carbide adhesion layers. The TIMEs were implanted in the median and ulnar nerves of three trans-radial amputees for up to six months. Here, we present the characterization of the electrical properties of the thin-film metallization as well as material status *post explantationem* for the first time. The TIMEs showed reliable performance in terms of eliciting sensation and stayed within the electrochemical safe limits maintaining a good working range with respect to amplitude modulation. After termination of the trials and explantation of the probes, no signs of corrosion or morphological change to the thin-film metallization was observed by means of electrochemical and optical analysis. Damage to the metallization was assigned exclusively to mechanical impacts during explantation and handling. The results indicate that thin-film metallization on polymer substrates is applicable in permanent implant system.

## 1. Introduction

Several research groups developed technological solutions to provide sensory feedback for bidirectional prosthetic limb control [1–5] to reduce rejection of prosthesis use [6,7], improve user’s quality of life and increase dexterity [1,2,8–12] as well as embodiment with a positive impact on that distressing condition named phantom limb pain (PLP) [11,13,–16]. In order to achieve optimal performance, peripheral nerve interfaces (PNIs) need to exhibit high spatial resolution, must show a limited foreign body reaction and must not exceed the electrochemical safe limits of stimulation [17]. Needless to say, that longevity and technical stability of the implanted devices are key to success in all studies, no matter which PNI design has been chosen. The ideal vision for the amputee would be a replacement of the missing limb with sensorized prosthesis, which is connected to such an interface and gives the patient sensory feedback [7,18]. Therefore, chronic stability of implanted PNIs is the ultimate prerequisite. By now, a limited amount of PNIs proved stability over a long period [3,19] like years.

A first good approach was introduced by Dhillon and co-workers [20]. They implanted longitudinal intrafascicular electrodes (LIFE) in human subjects sub-chronically. The drawback of conventional LIFEs is the low spatial stimulation selectivity with one contact per wire. To overcome this drawback, polyimide (PI) based thin-film longitudinal intrafascicular flexible electrodes (tf-LIFE) with a higher number of stimulation sites were developed and implanted sub-chronically into a human upper limb amputee [21]. Unfortunately, the maximum safe injectable charge limit was reached after two weeks and the stimulation had to be interrupted.

One of the simplest, very stable, common PNIs is the cuff electrode, which is implanted circumferentially around the targeted nerve [22,23]. Cuffs are very robust, but limited exciting the inner fascicles of a nerve and therefore limited in spatial selectivity [24]. To overcome this drawback Tyler & Durand developed the flat interface nerve electrode (FINE, manufactured with silicone rubber molding and spot welding) [25], which flattens and re-arranges the nerve for higher selectivity. Attention has to be payed to the force applied [26]. The FINE was successfully implanted chronically up to 3.3 years in the median and ulnar nerves of upper limb amputees [2,27,28].

Multi-channel microelectrode arrays (MEAs) were initially developed for intracortical application. Modified MEAs were sub-chronically tested in human median and ulnar nerve of upper limb amputees for four weeks [29–31]. Barrese et al. showed, that not only biological reasons like gliosis lead to failure of intracortical MEAs, but also material and mechanical causes [32]. They investigated the device stability on 78 chronically implanted devices in non-human primates via analysis of the signal recordings. 62 devices failed entirely and most of the failures occurred within the first year of implantation due to mechanical issues, particularly due to failures within the connector. A complete signal loss for all implants was predicted by about 8 years. Within a further study, Barrese et al. investigated eight MEAs implanted chronically in eight non-human primates using scanning electron microscopy (SEM). They observed several material failures. Corrosion of the platinum electrode tips occurred, as well as changes to the underlying silicon bulk material. Moreover, the protective insulation materials like parylene-c and silicone elastomer exhibited defects. The parylene-c was susceptible to crack formation and therewith delamination. Silicone elastomer used for device insulation delaminated at edges. The materials defects accumulated with time *in vivo*. These results showed that long-term stability of devices is an absolute requirement for chronic usage and patient safety of (active) implants.

In terms of spatial selectivity polyimide (PI) based transverse intrafascicular multichannel electrodes (TIME) [33] are very promising [24,34], as they are pulled through the nerve and can address as well the inner fascicles maintaining a steady contact-to-fascicle relationship in time. After verification *in vitro* and *in vivo* within small and large animal models [24,35,–37], PI based thin-film implants [21] and TIMEs showed promising sub-chronic clinical outcomes [1,38]. Use of iridium oxide as stimulation contact material with its high charge injection capacities kept the stimulation sites well in the chemically safe charge injection regime. Further analysis, development and improvement of the TIMEs in terms of material integrity by using adhesion promoters between thin-film metallization and the polymeric substrate [39], enabled chronic application in three upper limb amputees [10–12,16,40–42]. Within this study, we present the analysis of implants used in the mentioned clinical studies implanted for up to six months. Design changes, incorporation of adhesion layers and improvements of packaging and assembly increased stability and longevity, which is mandatory to head towards a fully implantable system for long-term use of years.

## 2. Materials and Methods

### 2.1. Design considerations of the thin-film electrodes

The design of the thin-film TIME passed through various changes in its applications since the very first idea in 2008 (**Figure 1**). From the early *in vitro* and acute *in vivo* experiments in rats with the TIME-1, we rapidly learned some basic pre-requisites of implantation and fixation which were implemented within the further small and large animal implants TIME-2 and TIME-3, respectively.

**Figure 1:**
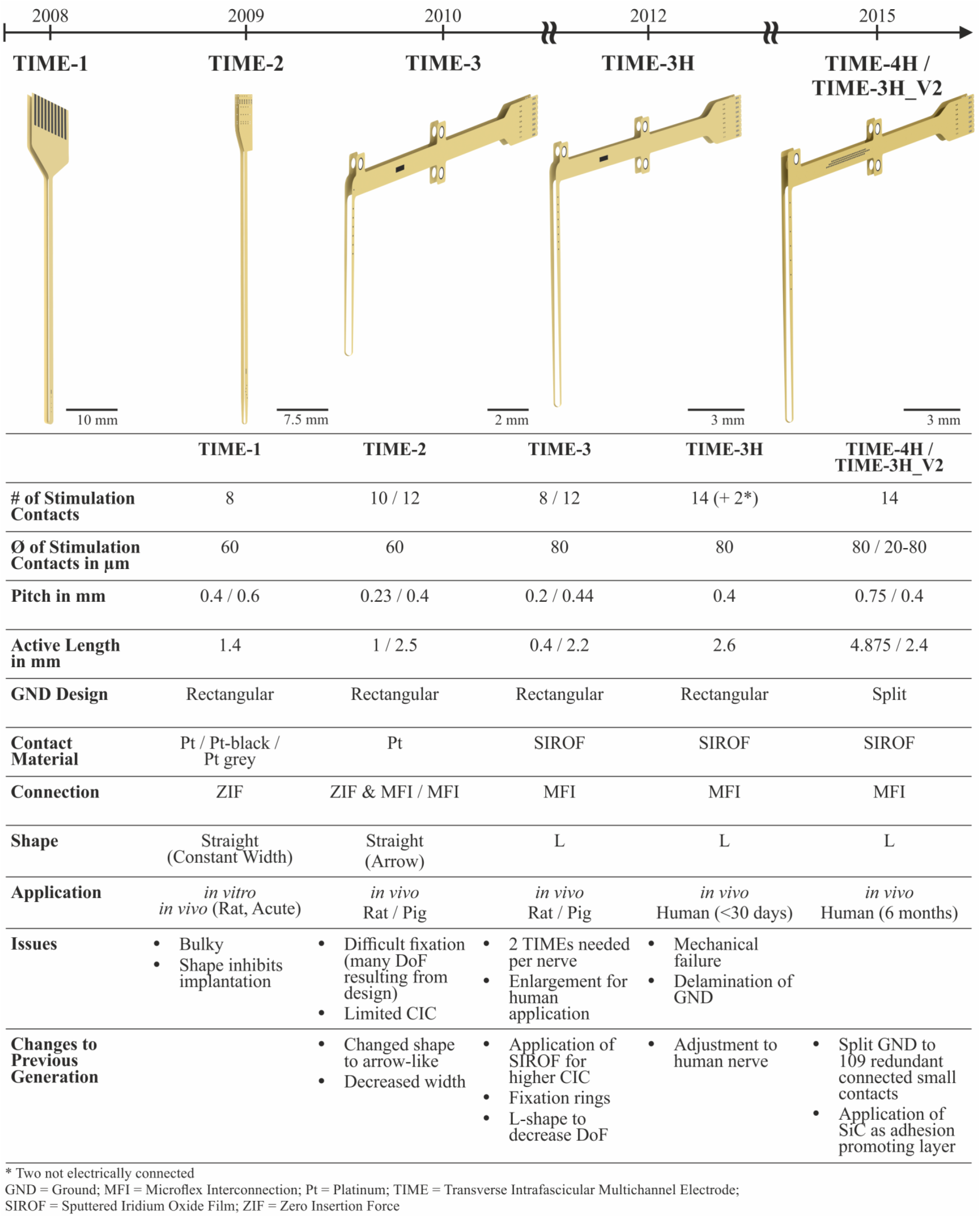
Design history of the TIME implants. TIME-2 was mostly used in small animal experiments, proving biocompatibility of the used materials [35] and showing advantages in the stimulation selectivity [24]. TIME-3 implants were used mainly in large nerves of pigs [36,37]. SIROF was applied as contact material to increase the charge injection capacity and thus increase the stimulation safety limits [17,45,46]. In a first-in-human study the TIME-3H showed excellent clinical outcomes [1,38], but as well weak points concerning the mechanical integrity, especially of the ground contact sites [39]. SiC was introduced as adhesion promoting layer and the large area ground contact site was split into 109 redundant connected small contact sites.

The implant version TIME-3H proved feasibility as an adequate peripheral nerve interface used for sensory feedback during the first sub-chronic human clinical trial [1,38]. The clinical outcome and the electrical stability of the implants were excellent. Due to explantation and subsequent handling of the thin-film electrodes, weak points were identified, especially concerning the mechanical integrity of the rectangular ground contact sites. As a first consequence, silicon carbide was introduced as an adhesion promoting layer between the PI and platinum. The new layer setup was tested and verified *in vitro* [43,44] and *in vivo* (small animal model) with regard to chronic application [39].

In order to improve the thin-film electrodes towards chronic implantation up to six months in humans, we made also some design considerations to lower intrinsic stress in the thin-film metallization, and thereby the risk of delamination, thus increasing the safety for the patient.

The TIME-3H [17,39] was consistently further developed up to the design freeze for the chronic human implantations, keeping fundamental design elements and changing details to improve the mechanical stability. The fundamental design elements were the two-dimensional U-shape for fabrication, which is transformed into a L-shape after folding during assembly. Details are described elsewhere [17,39]. Further fundamental design elements were realized with the thin-film sectioning. It consisted of three parts: the electrode, ribbon and transition part (**Figure 2**a). The electrode part contained the active strip with 14 stimulation sites incorporated (seven per side) and after folding a loop with an integrated surgical needle and thread. Positioning, pitch and size of the stimulation contact sites were depending on the thin-film version. The stimulation contact sites were labeled from the ribbon part towards the middle line L1 to L7 and R1 to R7 (**Figure 2**a). The ribbon part contained all tracks which led from the transition part to the stimulation contact sites and the large area ground sites, named L GND and R GND. The transition part had incorporated a structure to interconnect the thin-film electrode with a subsequent helically wound cable. The Microflex interconnection technique (MFI) [47] was used as a robust and reliable technique to assemble a thin-film ribbon with a long helically wound cable (**Figure 2**a).

**Figure 2:**
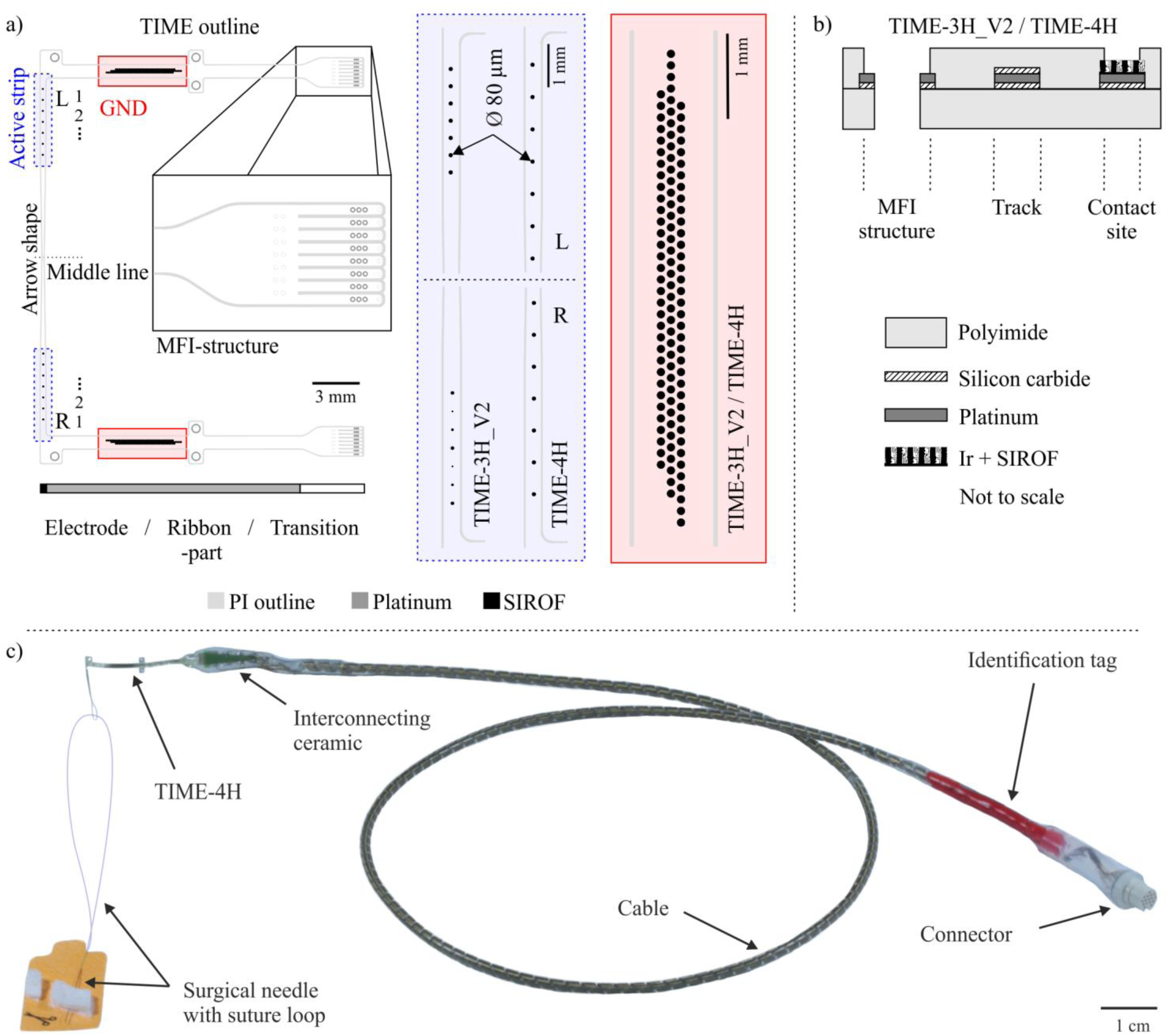
Overview of the different TIME designs. Two different designs for the active strips were realized (a, blue box). The ground contact site was split to 109 circular redundant connected contacts with a single diameter of 80 µm (a, red box). The layer setup of both designs was chosen identical (b). In the assembled TIME implant a surgical needle with a suture was incorporated within the thin-film loop (TIME-4H) (c). The polyimide based thin-film electrode was attached via the MFI technique to an interconnecting by use of screen-printing structured ceramic. To this in turn 16 helically wound MP35N-wires covered by a medical grade silicone rubber hose were soldered and terminated on the opposite side in a commercial connector. Colored silicone rubber and an identification tag made of laser structured platinum were used for distinct identification.

The focus of the new designs was placed on the large electrode area of the ground contact sites (see simulation sections). The ground contact sites are of fundamental importance, as a failure would lead to a total malfunction of the implant. The aim of the redesign was to prevent stress due to the large continuous area and rectangular edges, but to keep an area as large as possible to ensure an electrical closed-loop for stimulation. To preserve these boundary conditions, the rectangular ground site was split to 109 interconnected circular contacts with an exposed diameter of d = 80 µm (**Figure 2**a, red box) (diameter under polyimide d = 100 µm) of a single site. The contacts were arranged in a hexagonal assembly with a pitch of 125 µm in three rows. Eliminating large continuous areas and replacing highly stressed rectangular edges by round shapes should decrease the intrinsic stress of the thin-film metallization and thus decreasing the risk of delamination. A further advantage of the design, in case of partial damage, was the structural resilience of the design. The remaining ground contact sites would be still functional as every contact of the ground is electrically connected among each other.

Concerning the active strip, two new designs were developed, resulting in the TIME-3H_V2 and the TIME-4H. Both were manufactured with the split ground contact sites.

The TIME-3H_V2 was conceptualized in a way, that the left side of the electrode had integrated seven standard active sites with a diameter of 80 µm. However, the right side exhibited from the ribbon part downwards on the active strip decreasing diameters of the active sites. The diameter started at 60 µm and decreased with 20 µm steps down to a diameter of 20 µm and starting again at 60 µm. In folded state, the active sites were directly opposite to the left side. The active strip had a length of 2.4 mm, resulting in a contact site pitch of 0.4 mm.

The design of the active strip within the TIME-4H was developed in way to cope a larger area / diameter of the targeted nerve with a single electrode, as for this type of implant the surgical protocol was slightly different [48]. Considering that using directly opposite stimulation sites would mean one fascicle is addressed by two active sites, a shift of half the pitch (in this case 0.75 mm) of left and right side of the electrode was included to potentially address more fascicles with the same number of active sites. Otherwise the standard size for the active sites of 80 µm in diameter was used.

In both versions, platinum tracks and pads were sandwiched between polyimide as substrate and insulation layer (**Figure 2**b). Silicon carbide (SiC) was used as adhesion promoter between platinum and the PI substrate. Platinum tracks were completely surrounded by the SiC and polyimide (details below). At the active sites and ground contacts, the platinum layer was coated by iridium, subsequently covered by a sputtered iridium oxide film (SIROF) and opened via reactive ion etching (RIE). The MFI structure exhibited SiC only between platinum and the lower PI layer, in order to have bare platinum on the MFI ring after RIE opening for mechanical adhesion within the assembly procedure.

The detailed implantation procedure is described elsewhere [1,17]. In general, the front part of the TIME design (active part) is pulled with the incorporated needle and suture during the implantation through the nerve. The active sites for stimulation have to lay inside the nerve. The fixation flaps and the loop are sutured to the surrounding tissue, for the purpose of avoiding movements of the electrode during daily life.

### 2.2. Simulation of mechanical stress

Thermo-mechanical intrinsic stress was simulated using COMSOL Multiphysics^®^ (version 5.3, COMSOL Inc., Burlington, MA, USA). If not otherwise stated, standard parameters given by the software were applied.

Two concepts of the large area ground contacts were designed. First, a rectangular design used for the TIME-3H [17,39], i. e. with an exposed metallization area of 1 mm x 0.25 mm (A = 0.25 mm^2^) and second, a split ground with 109 circular exposed metallization contacts with a diameter of d = 0.08 mm (A = 0.55 mm^2^). For simplification, a polyimide-platinum sandwich was simulated without adhesion layers and without SIROF. Platinum is the main conductive material, representing the tracks, MFI structures and the base of the stimulation and contact sites. If failure occurs within the platinum structure, the whole device is prone to failure. Minimizing the intrinsic stress in the platinum layer is of highest relevance. SIROF however serves only as enhancement of the charge transfer to the tissue.

The geometries were designed in SolidWorks (Version 2014, Dassault Systemes Deutschland GmbH, Stuttgart, Germany) and imported in COMSOL Multiphysics^®^. Parameters of both materials, polyimide and platinum were acquired via the internal “MEMS” materials library (changed parameters are listed in **Table 1**). Several boundaries from the “Solid Mechanics” and “Heat Transfer in Solids” from the internal definitions were applied. An external temperature of 394.15 K as heat flux boundary was chosen according to the sterilization process performed after assembly (not 723.15 K imidizing temperature, since the metallization is homogeneously sandwiched between closed PI layers within this process step). A free tetrahedral mesh was used with custom changed mesh sizes.

**Table 1:** Changed parameters for the thermo-mechanical stress simulation of various ground contact sites.

| Material | Changed parameter | Value | Unit |
| --- | --- | --- | --- |
| Polyimide | Coefficient of thermal expansion | $30 \cdot 10^{-6}$ | K<sup>-1</sup> |
| Polyimide | Poisson's ratio | 0.34 | n. a. |
| Polyimide | Young's modulus | $8.83 \cdot 10^{-9}$ | Pa |

### 2.3. Cleanroom fabrication of thin-film electrodes

The microfabrication of the TIME thin-film electrodes was conducted in a cleanroom environment using standard photolithographic and MEMS processes (**Figure S1**). Details are described in the supplementary material and elsewhere [17,39]. All thin-film electrode designs described above feature polyimide (PI, U-Varnish S, UBE Industries, LTD., Tokyo, Japan) as substrate and insulation material. MEMS processes utilized were plasma-enhanced chemical vapor deposition of SiC (PECVD, PC310 reactor by SPS Process Technology Systems Inc, San Jose, CA, USA), evaporation of Pt (Leybold Univex 500, Leybold Vacuum GmbH, Cologne, Germany) and sputter deposition of Ir and IrO_x_ (Leybold Univex 500, Leybold Vacuum GmbH, Cologne, Germany). After the last fabrication step, the thin-film electrodes were pulled off the silicon wafer with a pair of forceps (**Figure S1**g) for assembly of the implants.

### 2.4. Chronic human implant

The TIME-3H_V2 and TIME-4H implants (called TIME implants if specification not relevant) were assembled out of four sub-modules (**Figure 2**c; for details see [17]). First, the thin-film part containing the contact sites for stimulation and the ground contacts to close the electrical circuit. Second, a screen-printed interconnecting ceramic to mechanically and electrically connect the thin-film electrode to the third part, a 40 cm long cable. The fourth module was a commercially available connector (NCP-16-DD, Omnetics Connector Corporation, Minneapolis, USA).

The fabrication procedure was similar to the fabrication procedure of the TIME-3H (details in [17,39]) with little adjustments due to chronic application in human (details in supplementary materials).

After some connector issues, with regard to strain relief during daily handling, loosing channels within the first two patients [10] the connector assembly was strengthened using a protective rubber hose (NuSil MED-4750, Freudenberg Medical Europe GmbH, Kaiserslautern, Germany) at the wire-connector transition (**Figure 2**c) for patient 3.

The TIME implants were fabricated within the fully ISO 13485 certified Laboratory for Biomedical Microtechnology of the Albert-Ludwig-University of Freiburg, Freiburg, Germany. Before implantation, each TIME was hydrated in order to increase the charge injection capacity [49], characterized and tested on functionality. Afterwards, it was washed, wrapped in sterile bags, labelled and steam sterilized at 121 °C and 2 bars for 21 minutes. The stimulation and ground contact sites of all delivered implants were fully functional.

Within clinical trials, MRI investigations are of interest and therefore the state of MRI compatibility of implants is relevant [50]. Medical devices can be declared according to the ASTM standard F2503 MR safe, MR conditional and MR unsafe [51]. We have used the F2503-08 version while the F2503-13 was released after our examinations had been finished. Investigation according to the relevant standards (ASTM F2052-14, F2213-06 & F2182-11a) have demonstrated that the TIME-4H implants (in clinically relevant position and orientation according to manufacturer specification) can be considered “MR conditional”. Meaning, a patient with these devices can be safely scanned in a MR system meeting the following conditions; 1) static magnetic field of 3 T, with 2) maximum spatial field gradient of 25700 Gcm^-1^ (257 Tm^-1^) and 3) maximum force product of 441 T^2^m^-1^.

Under the scan conditions defined above, the TIME-4H electrode is expected to produce a maximum temperature rise of < 0.5 °C (2.3 Wkg^-1^, 3 T) RF-related (switched gradient during imaging) temperature increase with a background temperature increase of < 0.1 °C (2.3 Wkg^-1^, 3 T).

### 2.5. Electrochemical electrode characterization

In order to compare the electrochemical properties of the TIMEs before implantation and after explantation the devices underwent a characterization with electrochemical impedance spectroscopy (EIS) *in vitro*. The characterization was carried out with a frequency analyzer and a potentiostat (SI 1260 & SI 1287, Solartron Analytical, Farnborough, UK) in phosphate-buffered saline (PBS) solution with a voltage amplitude of 10 mV and a frequency range of 100 kHz to 1 Hz. The three-electrode setup consisted of an Ag/AgCl reference, a large area platinum counter and the respective stimulation or ground contact sites of the TIME implant as working electrode. Beforehand, the SIROF was hydrated via cyclic voltammetry [49], using the identical setup as used within EIS. The voltage sweep was performed between - 0.6 V and 0.8 V, with 200 mV steps for 250 cycles.

Three neural stimulators: STIMEP (INRIA, Montpellier, France & Axonic, Vallauris, France), EARNEST (University of Cagliari, Italy) and the Grapevine Neural Interface System (Neural Interface Processor, 512 Channels of Potential, Ripple LLC, Salt Lake City, UT, USA) were used to inject current into the nerves of the subjects by means of TIME implants (details are provided in the supplementary materials). For the characterization of the electrode-nerve impedance *in vivo* and the subjects’ response to intraneural stimulation we delivered trains of cathodic-first, biphasic and symmetric square-shaped, current-controlled stimulation pulses of variable intensity, duration, and frequency, through a dedicated software controlling one of the before mentioned, external, electrical neural stimulators. The impedance was estimated as the ratio between the potential difference between the selected stimulation contact site and the ground contact sites (both, left and right one of the electrode) at the end of the cathodic pulse phase. The potential resulted from the average of four pulses (the first one of a five-pulse train was removed) with a current amplitude of 20 µA and a pulse width of 300 µs, repeated at a frequency of 1 Hz. While a safety margin has been applied in functional assessment in the patients [10], this paper uses 150kΩ as maximum impedance for the definition of electrically functional stimulation contact sites which corresponds to the maximum output swing of the stimulators.

Statistical methods were applied for further data analysis. The data acquired *in vivo* was longitudinal and unbalanced for each time point (weeks). Moreover, the data was not normal distributed and the variance was not homogeneous as well. Therefore, we applied a linear mixed effect model using the software R (version 3.5.2, The R Foundation for Statistical Computing, Vienna, Austria) and RStudio (Version 1.1.463, RStudio Inc., Boston, MA, USA), which is robust to non-normality and unbalanced data sets. Further detailed statistical analysis of the impedance data was performed elsewhere [10].

### 2.6. Optical analysis of explanted thin-film electrodes

The optical analysis of the stimulation and ground contact sites were done using light microscopy with polarization filters (Leica DM400M, Leica Microsystems GmbH, Wetzlar, Germany) to enhance visualization of surface irregularities and changes in adhesion (**Figure 3**, left column).

**Figure 3:**
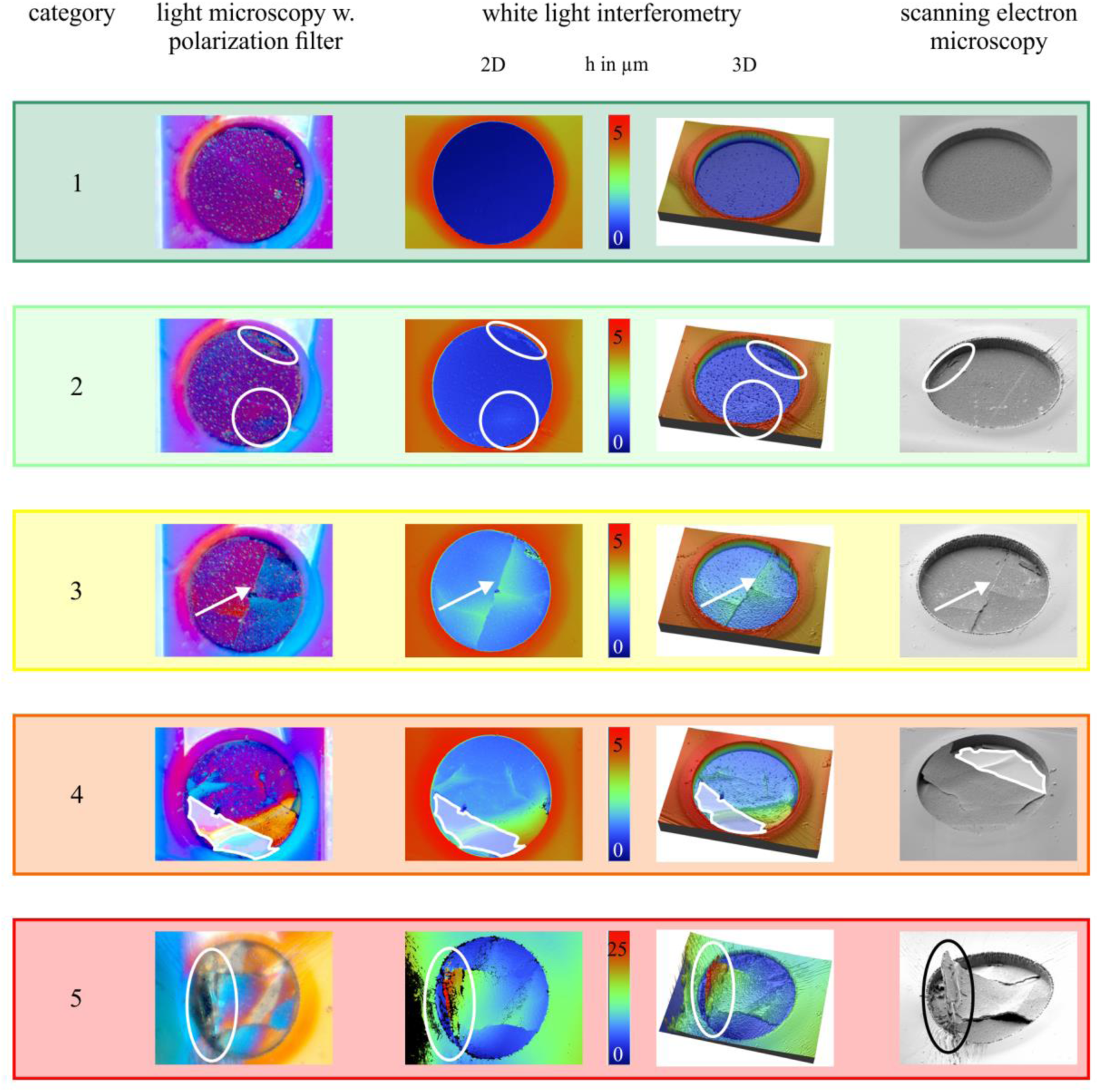
Acquisition samples of the optical thin-film analysis. Three methods were used: light microscopy with a polarization filter to make topographical irregularities visible, white light interferometry (WLI) for quantitative information in all three dimensions and scanning electron microscopy (SEM) in combination with focused ion beam (FIB) to gather high resolution overviews and information of the layer setup. Five categories were defined for the state of metallization integrity. Category 1 represents no impact and category 5 the highest.

White light interferometry (WLI; Wyko NT9100, Veeco Instruments Inc., Plainview, NY, USA) was used to quantitatively investigate the topography in three dimensions (**Figure 3**, middle columns).

In accordance to analyze the properties of the layer setup, scanning electron microscopy (SEM) in combination with focused ion beam (FIB; Zeiss Auriga 60, Carl Zeiss AG, Oberkochen, Germany) was utilized to gather high resolution overviews of the contact sites and to cut cross-sections into the thin-film metallization.

Five categories of stimulation contact site status were defined in order to obtain an objective classification (**Figure 3**). Category 1 represents a contact which can be compared to a pristine one, meaning from a material (not electrical) point of view, fully functional, with perfect adhesion and perfect surface integrity. Category 2 exhibits signs of delamination of ≤ 1 µm (distance between metal and underlying polyimide substrate) and / or (possible) light crack formation in the contact site coating. Delamination between 1 µm and 6 µm and crack formation occurs in category 3. Partial delamination of a metallization layer is categorized as cat. 4. The highest impact of destruction occurs in category 5 with heavy delamination of > 6 µm, disintegration of the metallization layers or compression or destruction of the PI substrate.

To gather compound overviews of the ground contact sites with a high degree of details, images with a table top SEM (Phenom Pro Desktop SEM, Thermo Fisher Scientific Phenom-World B.V., Eindhoven, Netherland) were acquired and highlighted with CorelDRAW X7 (Corel Graphics Suite X7, Corel GmbH, München, Germany).

### 2.7. Connector analysis

Issues with the connector assembly occurred as already mentioned above. During daily handling (connection and disconnection of the extracorporeal stimulator) many channels were lost. After scheduled termination of the clinical trial, the implants were retrieved. The connectors were analyzed with regard to the electrical state and the mechanical integrity. The electrical state was tested with a resistance meter (34401A 6 ½ Digit Multimeter, Agilent, Santa Clara, CA, USA) between different tapping points (connector – solder pads on interconnecting ceramic – wires anterior to interconnecting ceramic). Mechanical integrity at the wire-connector transition was investigated using µ-computer tomography (µ-CT; Nanotom m, GE Sensing & Inspection Technologies, Wunstorf, Deutschland) (**Figure 4**). Using this technique, x-ray images were acquired in all three dimensions throughout the wire-connector transition (**Figure 4**, right). In case of a ruptured wire from the connector there is no white dot representing the cross-section of a wire visible at the transition position (**Figure 4**, right, a). In the transition image in **Figure 4**, right, b) all 16 wires a visible.

**Figure 4.**
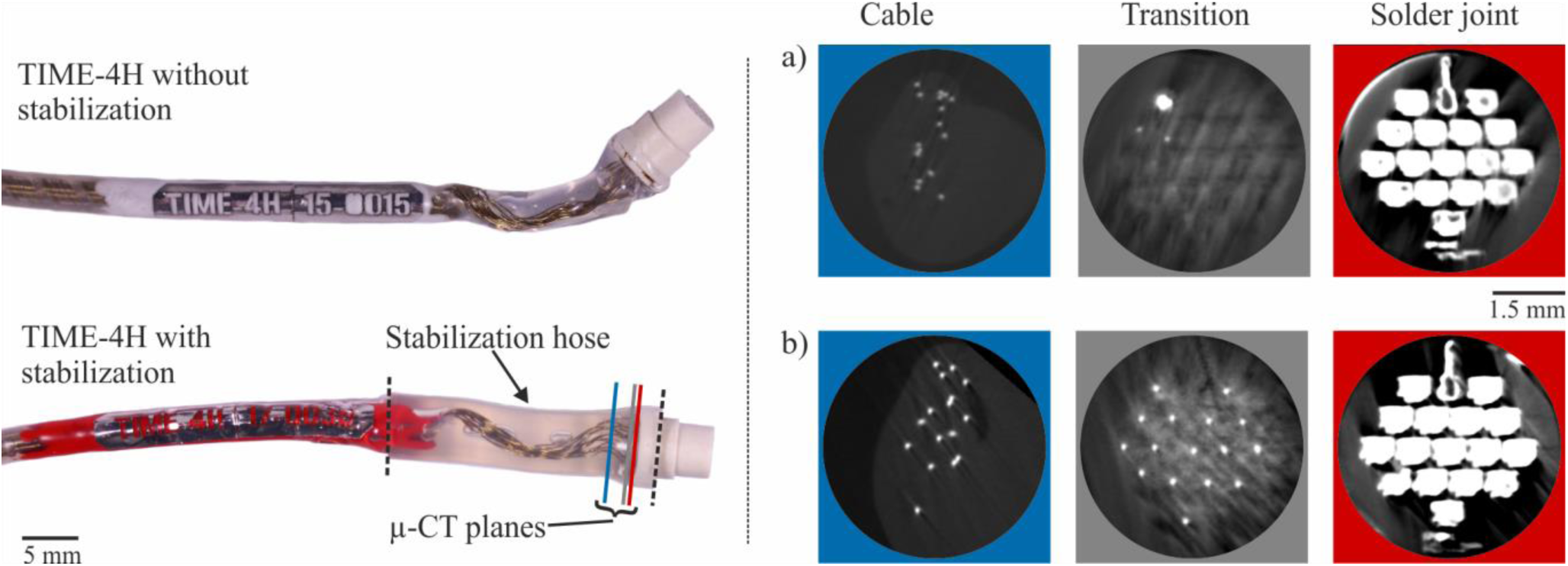
Analysis and stabilization of the wire-connector transition. On the left side, the upper connector part of the TIME implant has no stabilization incorporated as used for patient 1 and 2. Below, a stabilization hose was added to the transition between wires and connector for strain relief. The µ-CT planes for integrity analysis were labeled in the lower connector. On the right side, µ-CT images are representing a connector with ruptured wires (a) and mechanically integer (b) at transition.

## 3. Results

### 3.1. Simulation

The simulation of thermally induced stress due to the sterilization process was run with two different design approaches, a rectangular large area ground contact and a split ground with 109 circular interconnected contact sites (**Figure 5**).

**Figure 5:**
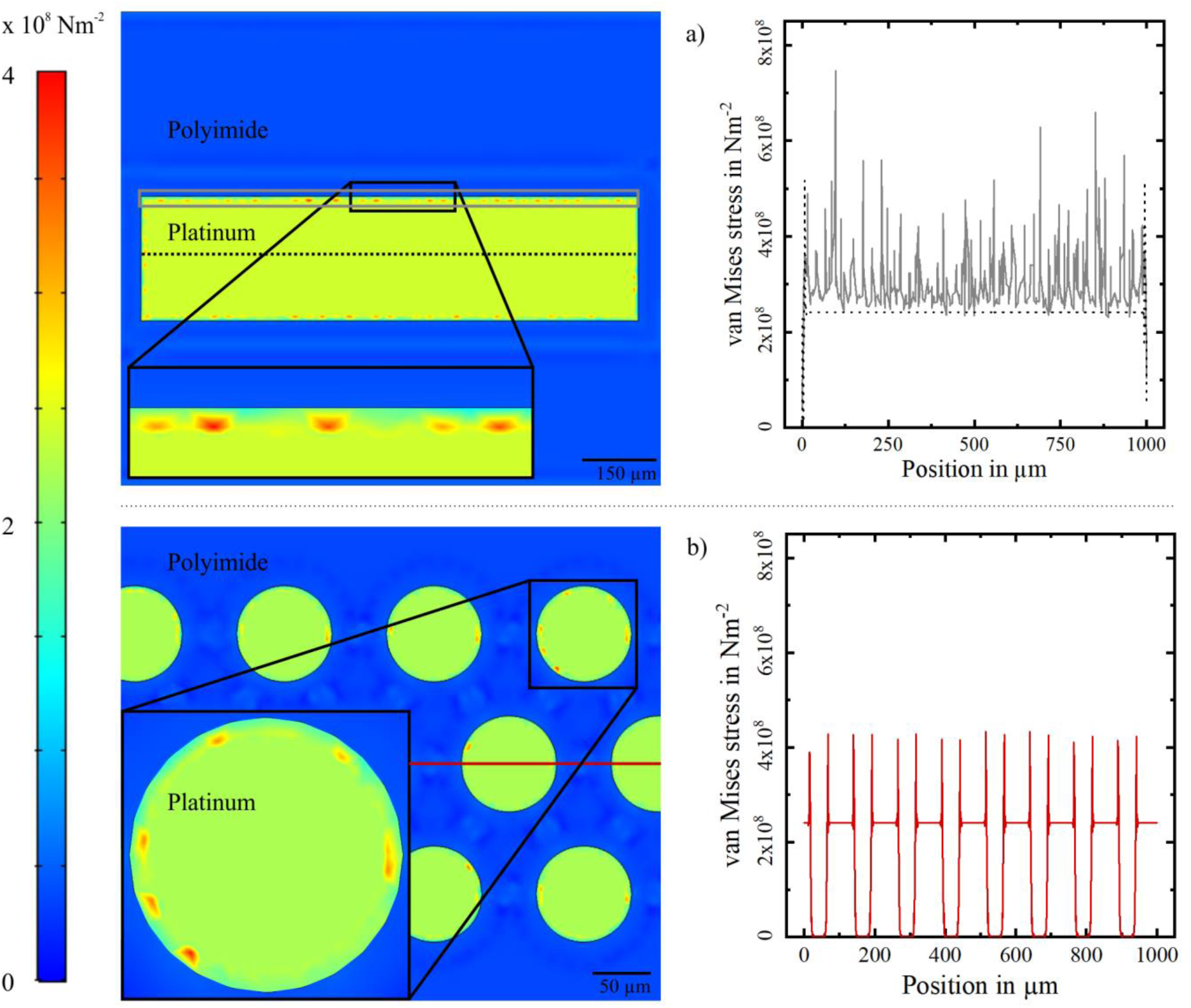
Simulation of thermal induced stress of different ground contact approaches. The rectangular ground contact site (a, 1 mm x 0.25 mm) was used on the TIME-3H within a sub-chronical trial [Cvancara2019]. Simulated stress values were acquired close to the edge of the PI (grey line) and in the middle of the contact (dashed line). The split ground with 109 single redundant connected contact sites (b, diameter of single contact 80 µm) was designed for chronic application in the TIME-3H_V2 and TIME-4H implants. Simulated stress values were acquired in the middle of the device on a length of 1 mm.

The van Mises stress level was indicated with a color range from low stress in blue to high stress in red. Generic, the stress was calculated in the middle of the rectangular ground contact superficial of the thin-film metallization within a low stress area (indicated by the dashed blackline, **Figure 5**a) with a length of 1 mm and with a distance of 8 µm from the edge of the polyimide in a high stress area (indicated by the grey box, **Figure 5**a). In the low stress segment, the stress did not exceed 2.4 x 10^8^ Nm^-2^, except at the edges (Position 0 µm and 1000 µm). In the high stress segment, the simulated van Mises stress was constantly above 2.4 x 10^8^ Nm^-2^, with peaks up to 7.5 x 10^8^ Nm^-2^. The integrated van Mises stress over the length of 1000 µm resulted in 3.0 x 10^11^ Nm^-1^ for the high stress segment, and 2.4 x 10^11^ Nm^-1^ for the low stress segment.

For the split ground contact, the simulated van Mises stress was calculated along the indicated red line with a length of 1 mm (**Figure 5**b). Peaks with a maximum stress of 4.3 x 10^8^ Nm^-2^ occurred at the edges of the exposed single contacts. Between the exposed thin-film metallization thus where the metallization was sandwiched between two polyimide layers, the stress dropped entirely. At other exposed metallization surfaces the stress stayed constant like above at 2.4 x 10^8^ Nm^-2^. The integrated van Mises stress over the length of 1 mm resulted in 1.6 x 10^11^ Nm^-1^.

### 3.2. Electrochemical characterization

The performance of the TIME implants was measured in terms of impedance of each stimulation contact. The impedance was acquired weekly for every patient during the clinical trials (**Figure 6**). The impedances of the implants used for patient 1 were from the beginning close to 100 kΩ. With a moderate slope of 0.8 kΩ/week the impedances reached a plateau little higher than 100 kΩ already in week 5. After week 22 a strong decrease in impedance occurred, reaching a mean impedance of 73 kΩ ± 18 kΩ close to termination of the clinical trial. The mean impedance of the devices of patient 2 was in the first week after implantation at a very low level of 21 kΩ ± 26 kΩ. From week 11 on, the impedances stabilized on a plateau of about 100 kΩ, resulting in a slope of 2.8 kΩ/week. The mean impedance of the implants of patient 3 increased from 60 kΩ ±36 kΩ in week 1 moderately to 67 kΩ ±36 kΩ, at the end of the trial resulting in a slope of 0.8 kΩ/week. In contrast to patient 1, there was no plateau in the middle of the clinical trial.

**Figure 6:**
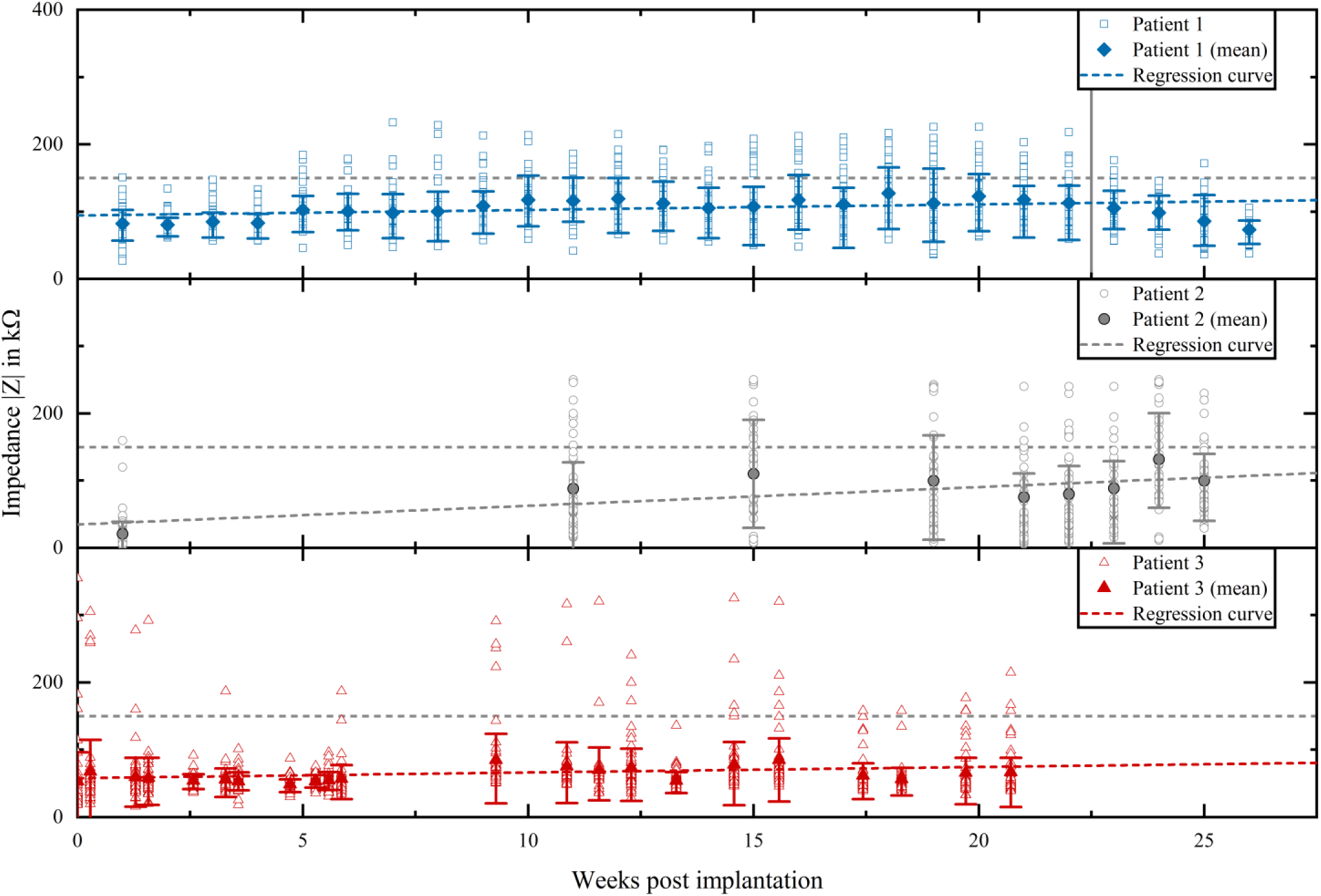
Evolution of impedance measured *in vivo* during the clinical trial of all patients. Transparent objects represent the single recorded values, filled objects represent the mean value of all contact on the day of measurement. The regression curve was evaluated using a linear mixed effect model. The slope for patient 1 was 0.8 kΩ/week, patient 2 2.8 kΩ/week and for patient 3 0.8 kΩ/week.

Prevention of nerve damage was of highest priority during the explantation procedure. Therefore, all cables were cut and the connector was removed before the explantation of the intrafascicular device was accessed. The thin-film parts of the implants had to be removed from the nerve carefully not to harm the patient. During explantation and handling after explantation all contacts but one ground contact were electrically disconnected. This single right ground contact (serial number TIME-4H | 17-0058, patient 3, ulnar nerve, proximal position) was characterized after explantation via electrochemical impedance spectroscopy (EIS, **Figure 7**). The impedance at 1 kHz decreased from 508 Ω before implantation to 355 Ω after explantation. Subsequently, the implant was hydrated using cyclic voltammetry. Further EIS displayed an impedance of 354 Ω at 1 kHz. The cut-off frequency increased from 5.8 Hz before implantation to 11 Hz afterwards. The hydration led to a decrease to 5.4 Hz.

**Figure 7:**
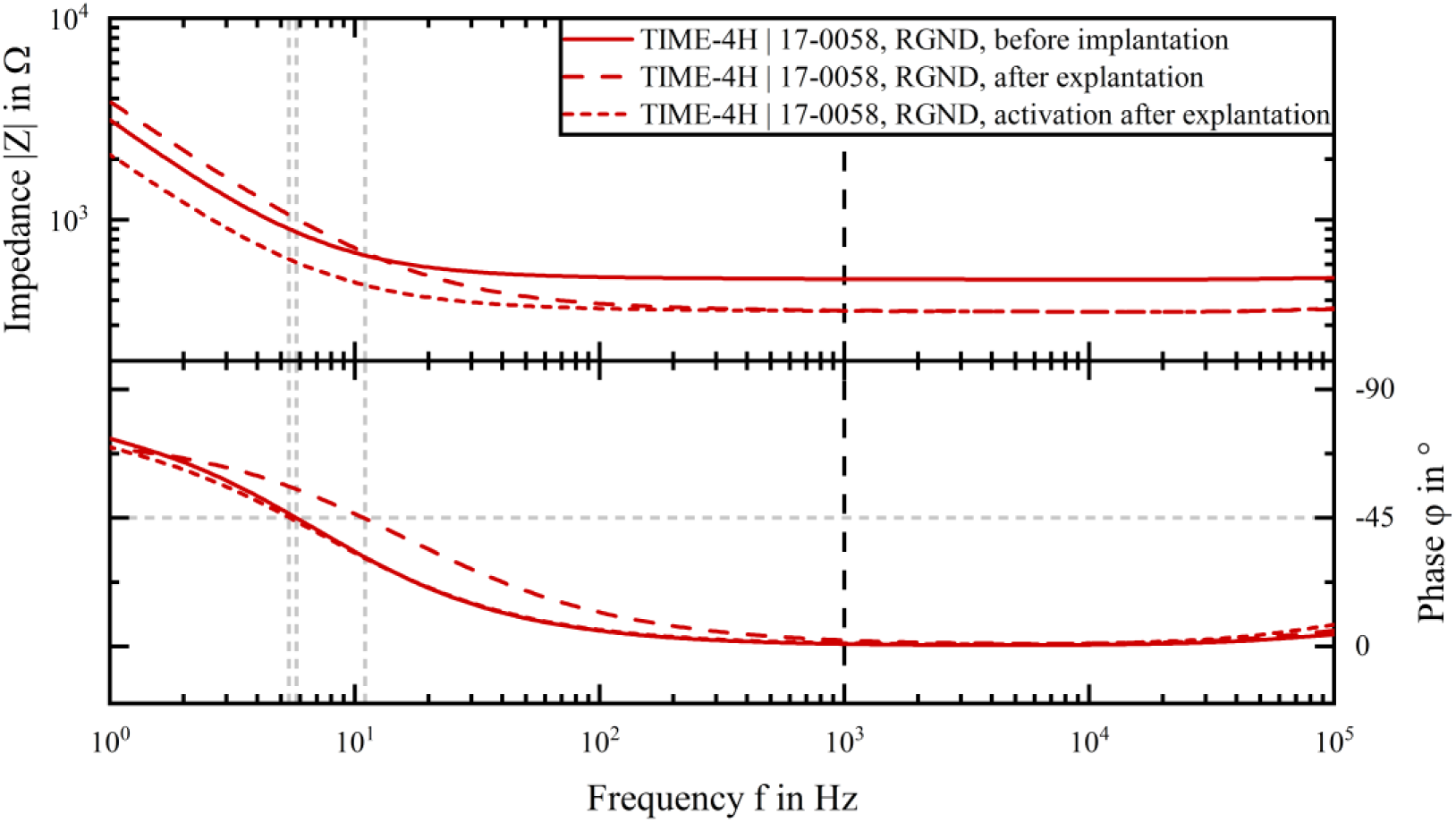
Characterization of the right ground contact of the implant with the serial number TIME-4H | 17-0058 before implantation and after explantation. Following a first characterization after explantation, the implant was hydrated and again characterized.

An electrically well-functioning stimulation contact site was defined having an estimated *in vivo* impedance of < 150 kΩ. With this definition, we analyzed the course of electrically functional contact sites in percentage over time in weeks post implantation (**Figure 8**, left). The dashed lines represented the real measured percentage, including fluctuation. The continuous lines ignored the fluctuation and visualized the highest effective percentage. The TIMEs used for patient 1 and 2 had a loss of electrically functional contacts from initial 98.2 % down to 73 %, respectively 68 % after 11 weeks. During the remaining course of the trial, a further decrease to a final 50 %, respectively 54 % could be observed. The percentage of electrically functional contacts of the devices implanted in patient 3 dropped from 96 % after week 1 to 82 % at the end of the clinical trial.

**Figure 8:**
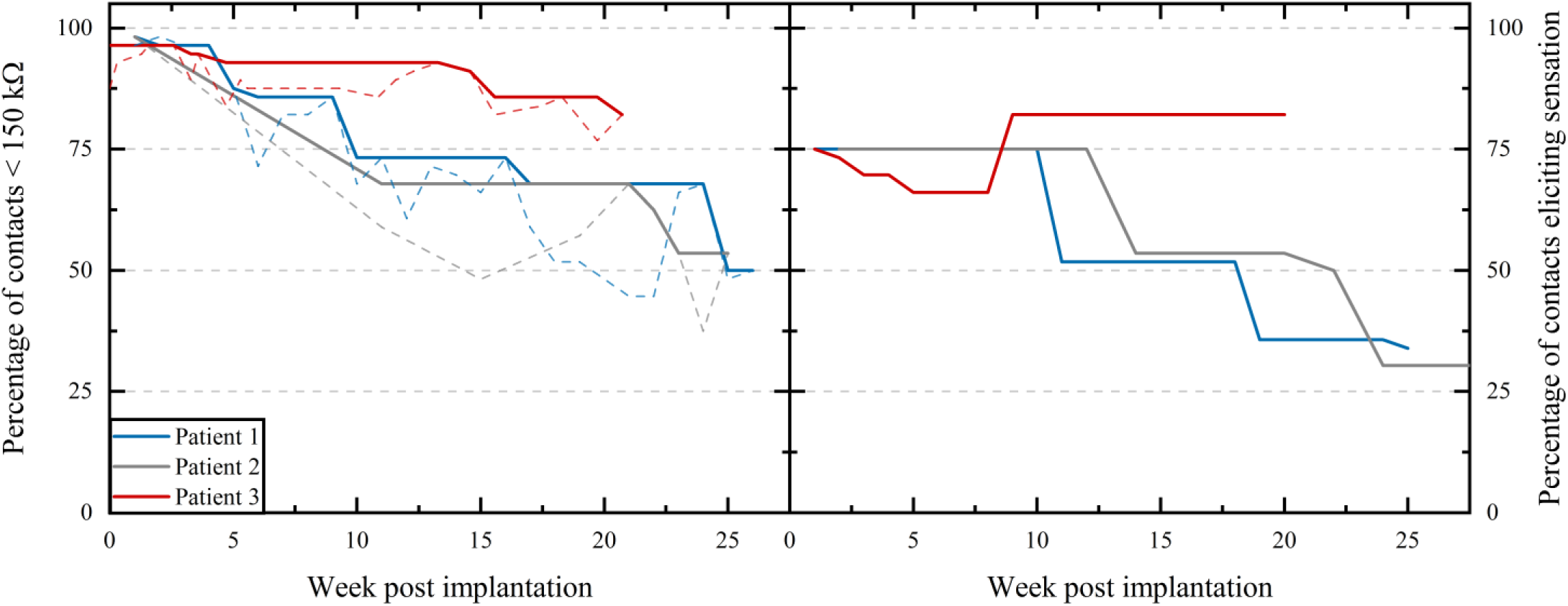
Kaplan-Meier-Plot comparing the percentage of electrically functional stimulation contacts (impedance < 150 kΩ) between the three chronic patients (left). Dashed lines display the real measured contact with fluctuation, within the continuous lines the fluctuation was filtered. Beyond, the percentage of contacts eliciting sensation reported by the patients was acquired over the time of implantation.

Next to the objective acquisition of electrically functional contacts, a subjective assessment of stimulation contact eliciting sensation was performed (**Figure 8**, right). Patient 1 felt until week 10 75 % of the contacts eliciting sensation, patient 2 until week 12. For both in the course of the clinical trial the percentage dropped to 34 % and 30 % respectively. Patient 3 communicated in the beginning (also starting at 75 %) a drop, with a stabilization after week 9 at 82 %, persistent until the end of the clinical trial.

### 3.3. Optical analysis of explanted thin-film electrodes

The impedances of the implanted TIMEs were recorded until the last day of the clinical trial, prior to explantation. Following explantation, the implants were optically analyzed in detail after thorough cleaning. The available data from the optical analysis (111 stimulation contact sites out of 168 implanted) were compared to the recorded impedance data close to explantation. The impedances of the stimulation contact sites were linked with the optically analyzed status to five categories (**Figure 9**). The categories correspond to those in **Figure 3**.

**Figure 9:**
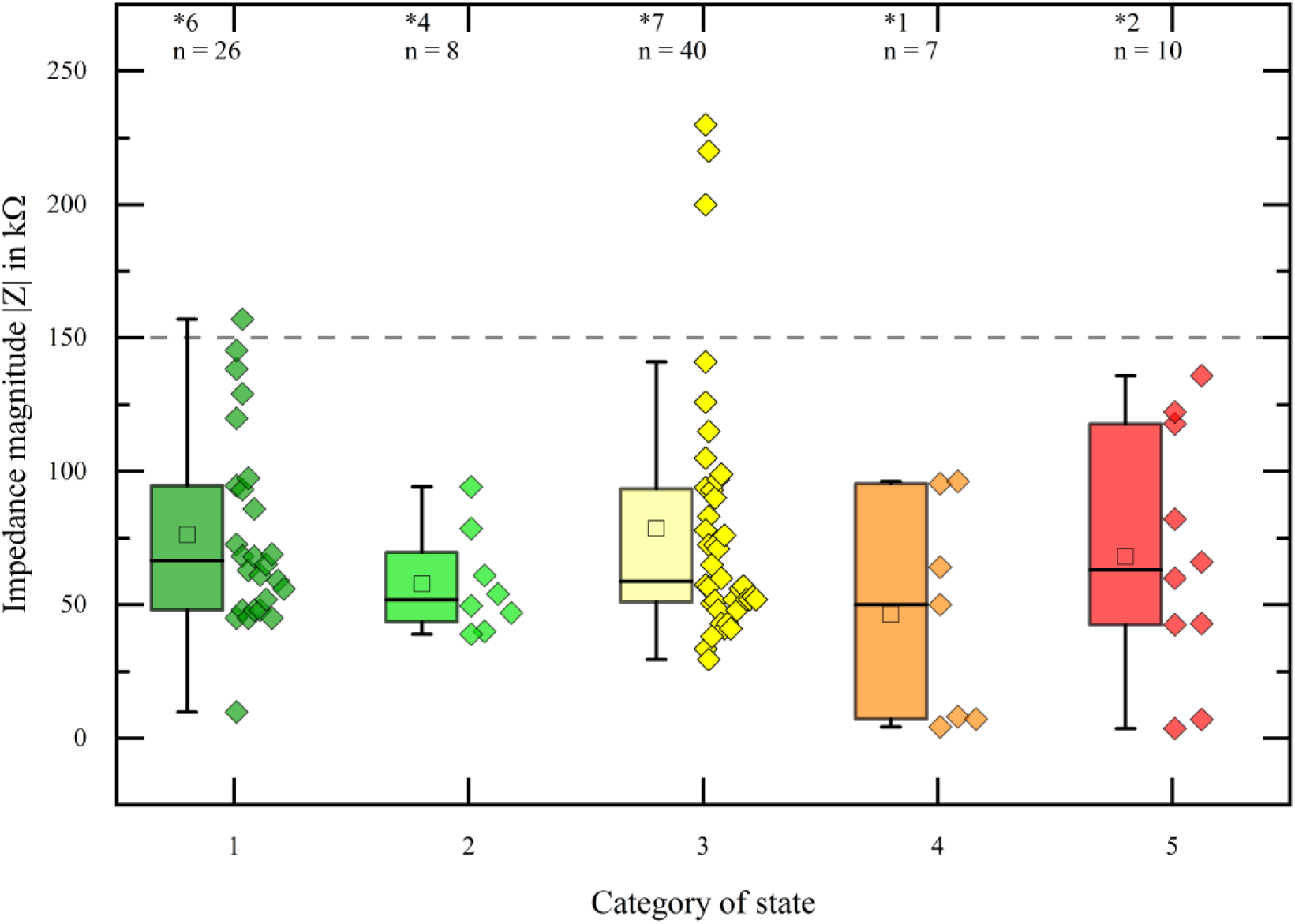
Distribution of impedances measured on the last day of use within the defined states of the metallization integrity. Box plots corresponding to the categories containing the mean (small box), median (bar) and the quartiles were generated. The numbers marked with a star represent not available impedance, n corresponds to the available impedances used for analysis. The impedance magnitude of 150 kΩ was defined as the limit of electrical functionality. There was no statistically significant difference between the impedances of the categories at the α-level of 0.05 (Kruskal-Wallis).

Could a certain impedance range be assigned to a classification of mechanical damage? An optically and mechanically pristine contact site corresponding to category 1 should exhibit a low impedance (< 150 kΩ), whereas an entirely destroyed contact site with for example missing metallization and corresponding to category 5 should either exhibit a very high impedance or even not be measurable.

The median impedances of all categories were between 46 kΩ and 79 kΩ (**Table 2**), thus within the definition of an electrically functional stimulation contact site (< 150 kΩ). The same applied for the median values, which were even in a narrower band of 50 kΩ to 67 kΩ. The majority of the impedance with 82 % came under category 1 to 3. Only 7 %, respectively 11 % were related to category 4 and 5. In **Table 2** the data was summarized.

A non-parametric Kruskal-Wallis method was applied, to assess differences between the impedances of the categories. The categorized impedances displayed no statistically significant difference at the α-level of 0.05.

**Table 2:** Acquired impedance data from last day of the clinical trial, referred to the categorization of the optical analysis.

| Category | Mean imp.<br>in kΩ | Standard deviation<br>in kΩ | Median imp.<br>in kΩ | No. of imp. | Available imp.<br>in % | No. of not<br>available imp. |
| --- | --- | --- | --- | --- | --- | --- |
| 1 | 76 | ± 35 | 67 | 26 | 29 | 6 |
| 2 | 58 | ± 18 | 52 | 8 | 9 | 4 |
| 3 | 79 | ± 47 | 59 | 40 | 44 | 7 |
| 4 | 46 | ± 38 | 50 | 7 | 7 | 1 |
| 5 | 68 | ± 44 | 63 | 10 | 11 | 2 |

Detailed images of the stimulation contact sites (111 out 168 available) were acquired using SEM. Especially on contact sites classified in category 5, traces of external mechanical impacts could be detected (**Figure 10**). The failure causes were associated to misuse during handling. The PI thin-film in **Figure 10**a) was bended heavily alongside the dashed blue line, resulting in total destruction of the metallization within the contact site. Groove marks in the PI resulting from the bending were highlighted with blue elliptic markers. A further mechanical impact originated from an object, which scratched the surface of the PI substrate and the thin-film metallization (**Figure 10**b), dashed red line). The detailed image revealed heavy damage to the thin-film metallization. In total 10 out of 12 optically analyzed category 5 stimulation contact sites exhibited distinct signs of external mechanical impacts. Ten exhibited impedances < 150 kΩ, while further two were not measurable. As already stated above, all implants were fully functional at the time point of delivery. A stimulation contact site from category 1 did not show any negative optical effect (**Figure 10**c). It exhibited on the last day of the clinical trial an estimated impedance of |Z| = 138.3 kΩ. A stimulation contact from category 2 exhibited slight crack formation without any negative effect on the stimulation parameters as it exhibited an impedance of |Z| = 94.2 kΩ (**Figure 10**d).

**Figure 10:**
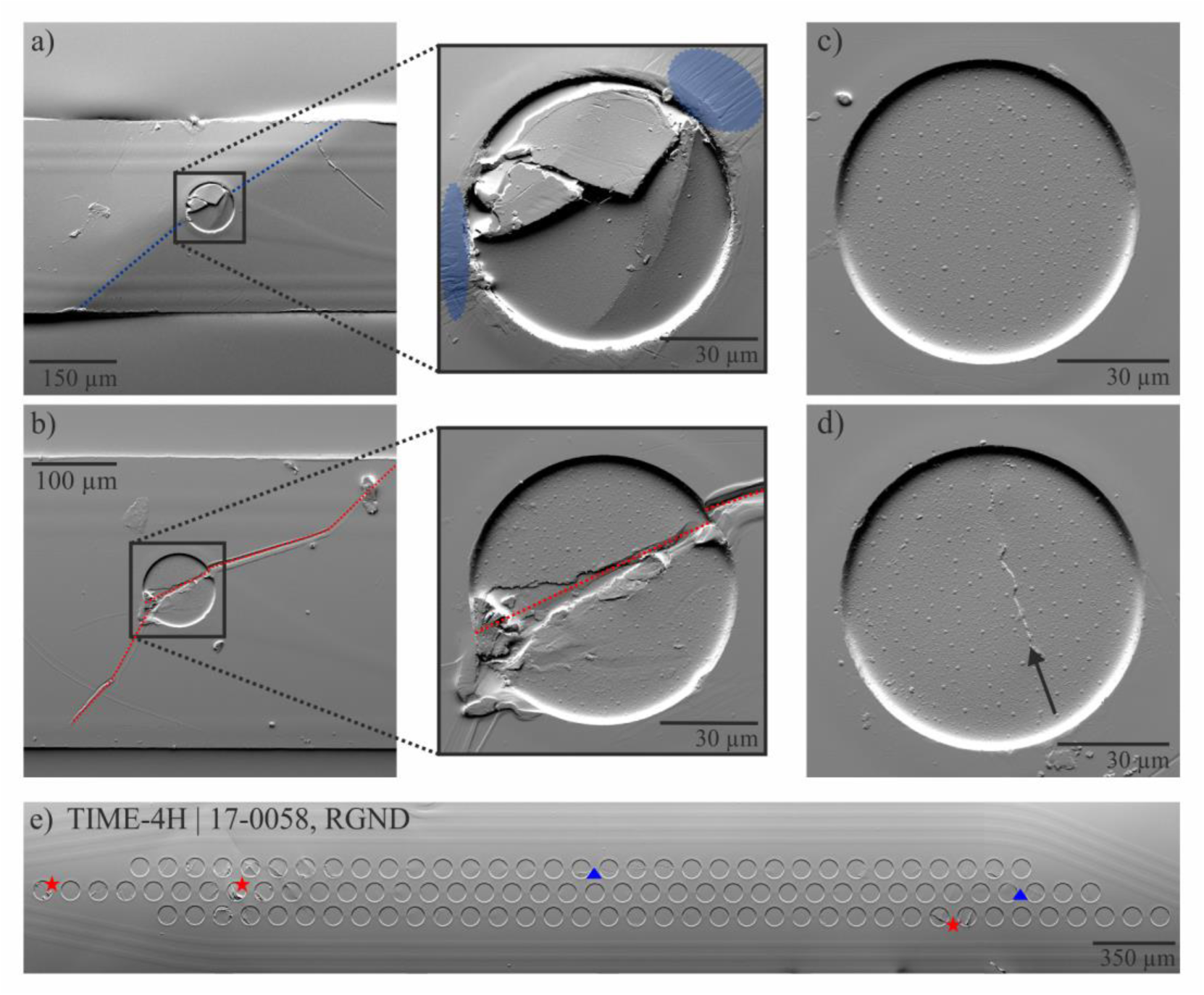
Detailed SEM images of stimulation contacts. Alongside the blue dashed line in a) the electrode was bend, which resulted in a kink. The detailed image shows within the blue highlighted areas groove marks from bending of PI. An object scratched the surface alongside the red dashed line in b). Metallization was heavily affected. The contact site in c) was classified in category 1 without any negative optical effect (|Z| = 138.3 kΩ). The contact in the detailed image d) exhibited little crack formation (arrow) and was classified in category 2 (|Z| = 94.2 kΩ). Overview of an explanted split ground used for 6 month during a human clinical trial in patient 3 (e). This ground contact was electrically functional after explantation. Red stars indicate crack formation with partial delamination. Blue triangles indicate crack formation.

Overview images of the ground contact sites were acquired after explantation, cleaning and when possible after electrochemical analysis. The implant with the serial number TIME-4H | 17-0058 was implanted in proximal position of the ulnar nerve of patient 3. After explantation it was electrochemically characterized using EIS (**Figure 7**) and exhibited an impedance of 354 Ω at 1 kHz. Thereafter, SEM images were acquired (**Figure 10**e). Approximately 50 % of the single ground contact sites displayed cracking or even partial delamination. Indicated by a red star, crack formation with partial delamination is shown. Slight crack formation is indicated by the blue triangles. More overview images of the ground contact sites can be found in the supplementary material. Predominantly, all via optical methods investigated ground contacts were mechanically intact, though they also displayed a certain amount of mechanical damage within the thin-film.

### 3.4. Connector analysis

The electrical analysis of the connectors (by measuring the resistance between connector and solder pad on the interconnecting ceramic) from patient 1 revealed that only two out of 64 channels (4 implants with 14 stimulation and 2 ground channels each) were electrically conductive (with a resistance of 178 Ω and 180 Ω respectively). Cutting the connector and measuring between the wires and the solder pads showed that all channels except one were electrically conductive and exhibited a resistance of 175 Ω ± 10 Ω. µ-CT analysis revealed 21 connected wires to the connector.

Electrical analysis of two connectors from patient 2 revealed that 22 out of 32 channels were electrically conductive. The mean resistance was at 169 Ω ± 40 Ω. According the µ-CT analysis 26 wires were connected to the connector, which corresponds to 81 %.

Within the retrieved implants from patient 3 62 out of 64 channels were still electrically conductive during electrical analysis. The channels exhibited a mean resistance of 171 Ω ± 13 Ω. µ-CT analysis revealed 64 wires were connected to the connector, which corresponds to 100 %. In **Table 3** all results of the electrical and µ-CT analysis of the connectors were summarized.

**Table 3:** Electrical and µ-CT analysis of the wire-connector transition part. The resistance was measured from the connector to the interconnecting ceramic. Only two connectors were available for analysis.

| Patient | Electrical analysis | | | μ-CT analysis | |
| --- | --- | --- | --- | --- | --- |
|  | Conductive channels |  | Mean resistance | Connected | wires |
| | # | % | R in $\Omega$ | # | % |
| 1 | 2 | 3 | $179 \pm 01$ | 21 | 33 |
| 2* | 22 | 69 | $169 \pm 40$ | 26 | 81 |
| 3 | 62 | 97 | $171 \pm 13$ | 64 | 100 |

## 4. Discussion

With the development of the TIME-3H using incorporated SIROF stimulation and ground contact sites we demonstrated the feasibility of PI based TIME implants as highly selective PNIs, capable providing for 30 days sophisticated sensory feedback to upper limb amputees [1,38]. After termination of the clinical trial and explantation of the devices, week points were identified, solved by introduction of adhesion promoting layers and validated in a small animal model [39]. Consequently, following questions arose: Are neural PI based thin-film electrodes capable being implanted for long-term use and stay mechanical of integrity? Will the impedances be stable during a chronic implantation and will the devices elicit stable sensations? Newly developed generations of TIME implants were thoroughly analyzed after six-month human clinical trials to answer these question and in order to identify problems and to introduce directly measures for following patients.

### 4.1. Simulation of a New Ground Contact Design and Verification *in vivo*

In order to decrease intrinsic stress in the metallization layer of the rectangular large area ground contact sites, we split it into 109 hexagonally arranged circular contact sites, which were redundantly connected between the PI layers. Stress levels were comparable to calculated ones in a similar work of Ordonez [52]. At the revealed metallization the stress levels were increased in both types of ground contact sites as there is no upper PI layer fixing the metallization. Near the PI edges intrinsic stress was up to three times higher than in the low stress middle of the contact sites. Particularly for the rectangular large area ground contact of the TIME-3H this led to a continuous high stress over the whole length of the contact near the edge. This is a potential risk for delamination, which was also observed by us in an earlier study [39]. The split ground contact exhibited similar peak stress levels, but very local, which was confirmed by the integration of the intrinsic stress over the same length.

Optical analysis post explantation showed, that the state of the split ground contacts was massively improved compared to the rectangular ground contacts used sub-chronically [39]. Although some single contacts of the grounds exhibited slight crack formation and / or delamination and mechanical impact like deformation caused by surgical forceps, this had no influence on the electrochemical characteristics of the ground contact sites, as all 109 contacts were redundant connected to each other.

### 4.2. Electrical characterization *in vivo*

Throughout the course of the six-month clinical trial with patient 1 we observed impedances around 100 kΩ. The percentage of electrically working stimulation contacts dropped to a level of 50 %. After week 22 we observed by pushing the cable of the implants towards the connector a regaining of the electrical contact, which also had an influence on the impedance slope during the clinical trial. Due to frequent connection and disconnection of the implants to the extracorporeal stimulator, the implants failed on the transition part between the wires of the cable and the connector. This was confirmed by the post-explantation electrical and µ-CT analysis. The differences between the electrical and µ-CT analysis can be explained by limitation of the µ-CT resolution of 25 µm. Similar lead breakage failures were described by other groups [32,53].

The connector problem was discovered and measures (stabilization and strain relief of the wire-connector transition) could be applied during respectively after patient 2. Therefore, we observed for patient 2 a similar failure rate compared to patient 1 including high impedance slopes as well. However, the modified wire-connector transition applied for the devices in patient 3 improved the performance significantly. The mean impedance was throughout the course of the clinical trial even below 100 kΩ. The number of contacts eliciting sensation stayed very stable, indicating that the thin-film has grown in the nerve very well, preventing displacement of the electrode [54]. The analysis of the wire-connector transition showed that 97 % of the channels were electrically conductive during and after the clinical trial. For the first time, we had the possibility to analyze a thin-film metallization electrically after chronic implantation post-explantation *in vitro*. The electrochemical characteristics remained identical to prior implantation. The shift to lower impedance could be explained by the cut cable due to safety reasons during explantation and therefore a reduction of the access resistance. After six months, no signs of corrosion could be determined within the SIROF contact layer.

### 4.3. Optical Analysis of the TIMEs from the chronic human clinical trial post-explantation

We classified the impedances recorded close to explantation of the TIMEs after optical analysis post-explantation in different categories. Category 1 represented stimulation contact sites with a pristine appearance, category 5 represented the highest impact of destruction with heavy delamination of > 6 µm, disintegration of the metallization layers or compression or destruction of the PI substrate. The categorized impedances displayed no statistically significant difference. The majority with 82 % of the impedances were in category 1-3 with no or only low mechanical damage. Only 11 % of the impedances were classified in category 5. However, externally caused mechanical damage by handling was identified in 83 % of these contacts as failure cause, indicated by groove marks following bending and scratches from heavy mechanical impact like scalpels (**Figure 10**) . This strengthens our presumptions made in our previous work, analyzing the implants from the first-in-human sub-chronical trial [39]. The thin-film parts of the implants were tightly encapsulated by surrounding tissue due to foreign body reaction [35]. Consequently, the thin-film experienced highest pulling and bending forces during explantation. The safety of the patient and the nerves must have always the highest priority, not the integrity of the TIMEs for potential *ex vivo* investigations. Several cuts had to be made to retrieve the implants in pieces, thus the evaluation had to be performed on fragments of the devices. The pulling and bending forces while explantation contributed to the observed crack formation and delamination of the metallization. Nevertheless, no morphological changes of the SIROF could be observed within all categories, indicating SIROF is an electrochemically stable material interfacing neural tissue.

## 5. Conclusion

We developed within this study a new generation of TIMEs for chronic – up to six months – micro-stimulation of median and ulnar nerves in upper limb amputees. Simulation of different ground contact designs helped us to lower intrinsic stress in the thin-film metallization of the implants, which was confirmed by optical analysis of the contact sites post-explantation. Electrical and µ-CT analysis of the implants used for patient 1 and 2 revealed, that the wire-connector transition was a critical location for failure, similar to previous chronic animal and human studies of other groups [32,53]. After stabilization of the transition and strain relief, the implants in patient 3 showed significant improvement in impedance stability and stimulation contacts eliciting sensation. Optical analysis strengthened the presumption of our previous work, that mechanical damage in stimulation contacts is caused by releasing the implants from tightly surrounded fibrotic tissue and not by electrical stimulation. The new ground contact proved to be stable throughout the chronic application. In this study we showed for the first time the feasibility of implanting thin-film electrodes for long-term applications. We are confident in longevity of thin-film electrodes and in the feasibility of polyimide-based permanent implants if the whole system design is long-term stable as such.

## Supporting information

Supplementary Material for Manuscript Cvancara - On the Reliability of Chronically Implanted Thin-Film Electrodes in Human Arm Nerves

## Acknowledgement

This work was supported by grants EPIONE (FP7-HEALTH-2013-INNOVATIO-1 602547) and NEBIAS (FP7/2007-2013-611687) from the European Commission (EC), and by the Bertarelli Foundation. The authors would like to thank the whole working group – especially those who are not on the list of authors – of the EPIONE and NEBIAS projects for their contributions and collaboration in the projects. Moreover, the authors would like to thank the KNMF at Karlsruhe Institute of Technology (especially Dr. Sabine Schlabach) for the granted working hours at the FIB (ID 2016-016013709).

## Funding Sources

This work was supported by the EU in its 7^th^ Framework Program [FP7-HEALTH-2013-INNOVATIO-1 602547]; and [FP7/2007-2013-611687].

## Author Contributions

P. Čvančara designed and fabricated the TIME-4H and performed the electrical analysis of the *in vivo* and *in vitro* measurements and the optical analysis of the implants. He transferred the outcome into the manuscript and was writing the manuscript.

Thomas Stieglitz was the co-designer of TIME-4H and TIME-3H_V2 and supervisor of this study, acted as scientific advisor and edited the manuscript.

G. Valle, F. Petrini, S. Raspopovic, I. Strauss, G. Granata, E. Fernandez, P. M. Rossini and S. Micera performed the human clinical trial and edited the manuscript.

M. Müller designed and fabricated the TIME-3H_V2 and edited the manuscript.

T. Guiho, A. Hiairrassary, J.-L. Divoux and D. Guiraud designed and manufactured the software and hardware of the stimulator STIMEP and edited the manuscript. T. Guiho, A. Hiairrassary and D. Guiraud acquired and preprocessed impedance data

M. Barbaro, designed and manufactured the software and hardware of the stimulator EARNEST and edited the manuscript.

K. Yoshida gave scientific advice in the clinical study and contributed in manuscript proof-reading.

W. Jensen gave scientific advice in the clinical study and contributed in manuscript proof-reading and editing.

## Disclosure

Declarations of interest: none.

## Figures

Color should be used for all figures in print.

