## Supplementary Material for Manuscript Cvancara - On the Reliability of Chronically Implanted Thin-Film Electrodes in Human Arm Nerves for "On the Reliability of Chronically Implanted Thin-Film Electrodes in Human Arm Nerves for Neuroprosthetic Applications"

[2] A!MD France

### 1. Cleanroom fabrication of the thin-film electrodes

All thin-film electrode designs described above feature polyimide (PI, U-Varnish S, UBE Industries, LTD., Tokyo, Japan) as substrate and insulation material. The first step was spin coating of 5 µm of PI on a 4”-silicon wafer and imidizing it at 450 °C under nitrogen atmosphere in a furnace (YES-459PB6-2PE-CP, Yield Engineering Systems Inc., San Jose, CA, USA), as shown in **Figure S1**a. Afterwards high-resolution image reversal resist (1.4 µm, AZ 5214E, MicroChemicals GmbH, Ulm, Germany) was applied via spin coating. Deposition of SiC as adhesion promoter (50 nm) was realized using plasma-enhanced chemical vapor deposition (PECVD, PC310 reactor by SPS Process Technology Systems Inc, San Jose, CA, USA). Subsequently, a 300 nm platinum layer was deposited via evaporation (Pt, Leybold Univex 500, Leybold Vacuum GmbH, Cologne, Germany) (**Figure S1**b). A lift-off step with acetone and isopropyl alcohol was performed to remove the resist with the excessive material. The next step was to pattern a further layer of image reversal resist, for the upper 40 nm SiC layer, which had to cover all metallization, for future contact with the top PI layer. (**Figure S1**c). After lift-off, a third layer of image reversal resist was applied to define the structures for the sputter desposited 100 nm iridium and following 800 nm sputtered iridium oxide film (SIROF) (Ir, Leybold Univex 500, Leybold Vacuum GmbH, Cologne, Germany) (**Figure S1**d). Excessive iridium and SIROF was removed by a lift-off step. The layer setup underwent an $O_2$-plasma activation step in a reactive ion etching chamber (RIE, RIE Multiplex, STS Surface Technology Systems plc, Newport, UK), followed by spin coating and imidization of the second 5 µm thick PI layer at 450 °C under nitrogen atmosphere (**Figure S1**e). Application of 30 µm thick positive resist (AZ 9260, MicroChemicals GmbH, Ulm, Germany) as etching mask was realized, followed by RIE in an oxygen plasma to open the perimeters and contact sites (**Figure S1**f). After the last fabrication step, the thin-film electrodes were pulled off the silicon wafer with a pair of forceps (**Figure S1**g) for assembly of the implants.

Both, the TIME-3H_V2 and TIME-4H thin-films used for the human clinical trials were investigated with special cytotoxicity samples [1] (run in parallel with the same process) according to the ISO 10993 for cytotoxicity testing. Direct contact and extract tests were performed compliant with the ISO 10993-5 with L929 mouse fibroblasts. Additionally, further direct contact and extract tests were performed with the human nerve cell line Kelly and the human muscle cell line A673. The samples passed the tests and no objections were claimed.

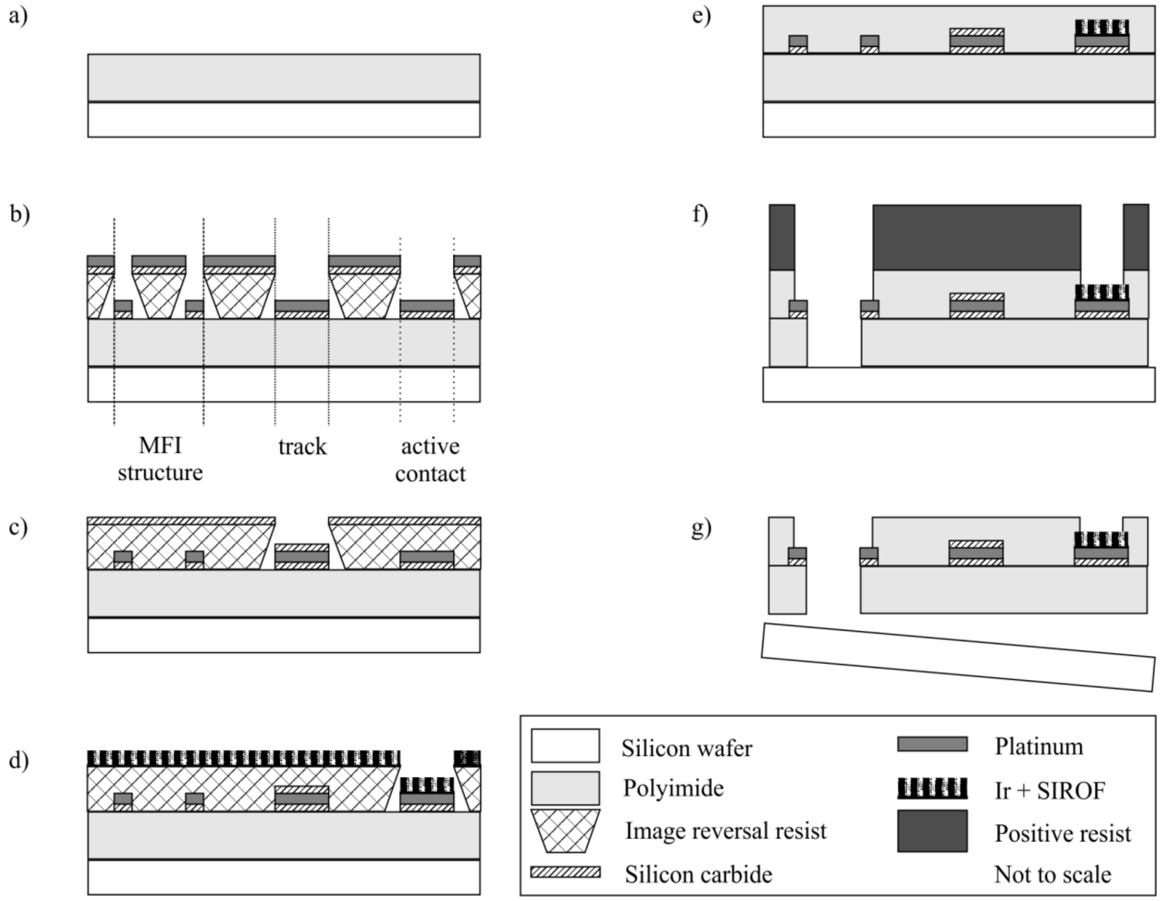


**Figure S1**: Fabrication process of TIME-3H_V2 and TIME-4H thin-film electrodes. After spin-coating and imidizing 5 µm PI (a), an image reversal resist was applied as structuring mask. Next, 50 nm SiC were applied via PECVD and 300 nm of platinum were evaporated (b). Following a new layer of image reversal resist, 40 nm SiC were created (c). After another image reversal resist, iridium and SIROF were sputter deposited on the active and ground contact sites (d). For electrical insulation a second layer of 5 µm PI was spin-coated and imidized (e), before opening the perimeters and the contact sites with RIE (f).

### 2. Assembly of chronic human implant

In a first step, 16 helically wound wires (MP35N with polyesterimide insulation) were inserted in a medical grade silicone rubber hose (NuSil MED-4750, Freudenberg Medical Europe

GmbH, Kaiserslautern, Germany) which was filled in turn with liquid silicone rubber (cured while assembly procedure). The wires of the 40 cm long cable were soldered into a circular ceramic-connector assembly (fabrication based on [2,3]) for electrical connection to an extracorporeal stimulator. The other end of the cable was soldered to two screen printed pre-structured interconnecting ceramics ($Al_2O_3$-96%-Rubalith 708S, A.L.L. Lasertechnik GmbH, Munich, Germany; 5837-G and 4771-P, ESL Europe, Agment Ltd., Reading, UK). The thin-film part was attached via the MFI technique to the interconnecting ceramics and stabilized with an epoxy (UHU Endfest Plus 300, Bolton Adhesives, Rotterdam, Netherlands). After folding the thin-film and bonding the ceramics backside to backside, a medical grade silicone rubber hose (NuSil MED-4750, Freudenberg Medical Europe GmbH, Kaiserslautern, Germany) was pulled over and filled with silicone adhesive (MED-2000, NuSil Technology LLS, Carpinteria, CA, USA). Colored medical grade silicone rubber (MED-2000 with MED-4800 color masterbatch, NuSil Technology LLS, Carpinteria, CA, USA) and an identification tag made of laser structured platinum was incorporated for a distinct identification near the connector.

### 3. Electrode characterization *in vitro*

Electrochemical characterization of the different TIME designs with their various stimulation contact sizes was performed using electrochemical impedance spectroscopy (EIS). The electrochemical properties were summarized in **Table S1** containing information about the magnitude of impedance $|Z|$ in kΩ and the phase shift $\varphi$ in ° both at 1 kHz for all contact sizes and types.

**Table S1**: Main electrochemical properties of various TIME designs, contact types and sizes. The data were acquired via EIS.

| | Type (diameter) | Impedance \|Z\| in kΩ @ 1 kHz | Phase φ in ° @ 1k Hz | Cut-off frequency $f_{cut}$ in Hz |
|---|---|---|---|---|
| Stimulation contacts | TIME-3H_V2, (80 µm) | 5.3 | -6.0 | 56.4 |
| | TIME-3H_V2, (60 µm) | 7.4 | -8.0 | 72.1 |
| | TIME-3H_V2, (40 µm) | 11.6 | -10.3 | 95.55 |
| | TIME-3H_V2, (20 µm) | 27.3 | -16.6 | 146.3 |
| | TIME-4H, (80 µm) | 5.7 | -9.8 | 79.8 |
| GND contacts | TIME-3H_V2 | 0.5 | -0.7 | 5.0 |
| | TIME-4H | 0.5 | -1.1 | 6.4 |
| | TIME-4H, RGND, before impl. | 0.5 | -0.8 | 5.8 |
| | TIME-4H, RGND, after expl. | 0.4 | -2.1 | 11.0 |
| | TIME-4H, RGND, re-hydrated | 0.4 | -1.0 | 5.4 |

### 4. Subject recruitment and surgical procedure

Three trans-radial amputees were included in the study [4]. Patient 1 was a 37-year-old male, left-handed, who had a traumatic trans-radial amputation of the distal two-thirds of the left forearm 2 years before the enrollment in the trial. Patient 2 was a 48-year-old female, right-handed, with a traumatic transradial amputation of the distal third of the left forearm which occurred 23 years before the enrolment. Patient 3 was a right-handed, 53-year-old female trans-radial (distal two-thirds of the left forearm) amputee. The amputation occurred in December 2015, following a traumatic accident at work.

Four TIME-4H were implanted in the median and ulnar nerves of patient 1 on November $14^{th}$ 2015, four TIME-3H_v2 in patient 2 on June $24^{th}$ 2016, and again four TIME-4H in patient 3 on June 24th 2017.

During general anesthesia, through a 15 cm-long skin incision on the left arm, the median and ulnar nerves were exposed to implant a proximal and a distal TIME in each nerve. The thin-film electrodes and a segment of their cables were placed through the long skin incision into proximity with the exposed nerve, two laterally and two medially to the center of the main surgical cut. The cable segments were located in subcutaneous pockets, externalized through four small skin incisions in order to be available for the transcutaneous connection with a neural stimulator. Then, using an operating microscope (Pentero-900, Carl Zeiss Meditec AG, Jena, Germany), the microelectrodes were implanted transversally within the nerve fascicles. Appropriate sutures and subcutaneous strain release loops secured the nerve implants. To verify the integrity and functionality of the implant, the impedance of the electrodes active sites was monitored intraoperatively. After 180 days, the microelectrodes were removed in accordance with the protocol and the obtained permissions.

Ethical approval was obtained from the Institutional Ethics Committees of Policlinic A. Gemelli-IRCCS at the Catholic University, Rome, Italy, where the surgery was performed. The protocol was also approved by the Italian Ministry of Health, division for experimental devices. All the subjects signed the informed consent.

Explanted devices were rinsed in ethanol and water. Afterwards, they were exposed to trypsin and again rinsed with water. Thin-film electrodes were prepared for optical analysis and cables were steam sterilized at 121 °C and 2 bar for 21 minutes

### 5. Neural stimulators

Three neural stimulators: STIMEP, EARNEST and the Grapevine Neural Interface System were used to inject current into the nerves of the subjects by means of TIME implants.

STIMEP (INRIA, Montpellier, France & Axonic, Vallauris, France) is a wearable neural stimulator based on a generic neural stimulation architecture [5,6]. It allows the control of 64 poles of implanted electrodes, divided in four ports of 16 poles each. Changing pulse-width, intensity as well as frequency, up to four TIMEs can be driven in real-time. Each port addressing each TIME includes 14 capacitive coupled active outputs (channels) and 2 non-capacitive coupled reference contacts. STIMEP further embeds safety procedures (charge limit) and impedance measurements. This stimulator was used for all the tests of patient 1 and partially patient 3 (impedance estimation).

EARNEST (University of Cagliari, Italy) is a prototypical, wearable, complete embedded platform for neural prosthetic applications [7]. It allows to simultaneously control and use up to 4 implantable multi-site (16 channels) electrodes, for a total of up to 64 stimulation- and recording-channels. The device, simultaneously, allows the recording of neural signals and as well as fully programmable electrical stimulation in terms of pulse-width, intensity and frequency. This stimulator was used with patient 2.

The Grapevine Neural Interface System (Neural Interface Processor, 512 Channels of Potential, Ripple LLC, Salt Lake City, UT, USA) is a commercial device that can be used for the recording of neurophysiological data and for delivering current-controlled stimulation through up to 32 high-impedance microelectrodes. Pulse-width, intensity and frequency were modifiable. The system was used on patient 3.

All stimulators were limited to the maximum safe charge injection capacity of $Q_{max,inj}$ = 2.3 $mCcm^{-2}$ of the iridium oxide contact sites.

### 6. Sensation characterization procedure

Regarding the evaluation of the patients percepts to the intraneural stimulation, we performed a sensation characterization procedure (or mapping) [4,8] in which the patient was connected to the neurostimulator using the transcutaneous cables of the implanted TIMEs. During this procedure, the stimulation was triggered by the experimenter. Trains of cathodic-first, biphasic and symmetric square-shaped pulses of increasing amplitude from 10 µA to 980 µA (steps of 10 µA), and fixed pulse width (chosen in the range 10 µs to 120 µs depending on the active site) and frequency (50 Hz as in our previous studies [4,9,10]) were injected. In each of these trains of increasing amplitude, the subjects had to press a button to stop the stimulation ramp when and whether they perceived a sensation. Then, they had to report the sensation properties (location, type, quality and intensity). When stimulating from an active site, the subject reported a reliable percept, then that active site was considered to be functional (in terms of sensory feedback).

At the end of this procedure, a map of sensation types, locations, extents and intensities (including the sensory threshold) referring to the correspondent active sites was obtained.

The sensation characterization was performed once per week or every two weeks, according to the availability of the subjects. In particular, patient 1 and 2 executed it weekly, while patient 3 performed them every two weeks.

### 7. Optical Analysis

In order to gather compound overviews of the ground contact sites with a high degree of details, images with a table top SEM (Phenom Pro Desktop SEM, Thermo Fisher Scientific Phenom-World B.V., Eindhoven, Netherland) were acquired and highlighted with CorelDRAW X7 (Corel Graphics Suite X7, Corel GmbH, München, Germany) (**Figure S2**). Patient 1 received the implants with the serial numbers “TIME-4H | 15-xxxx”, patient 2 “TIME-3H_V2 | 16-yyyy” and patient 3 “TIME-4H | 17-zzzz”.

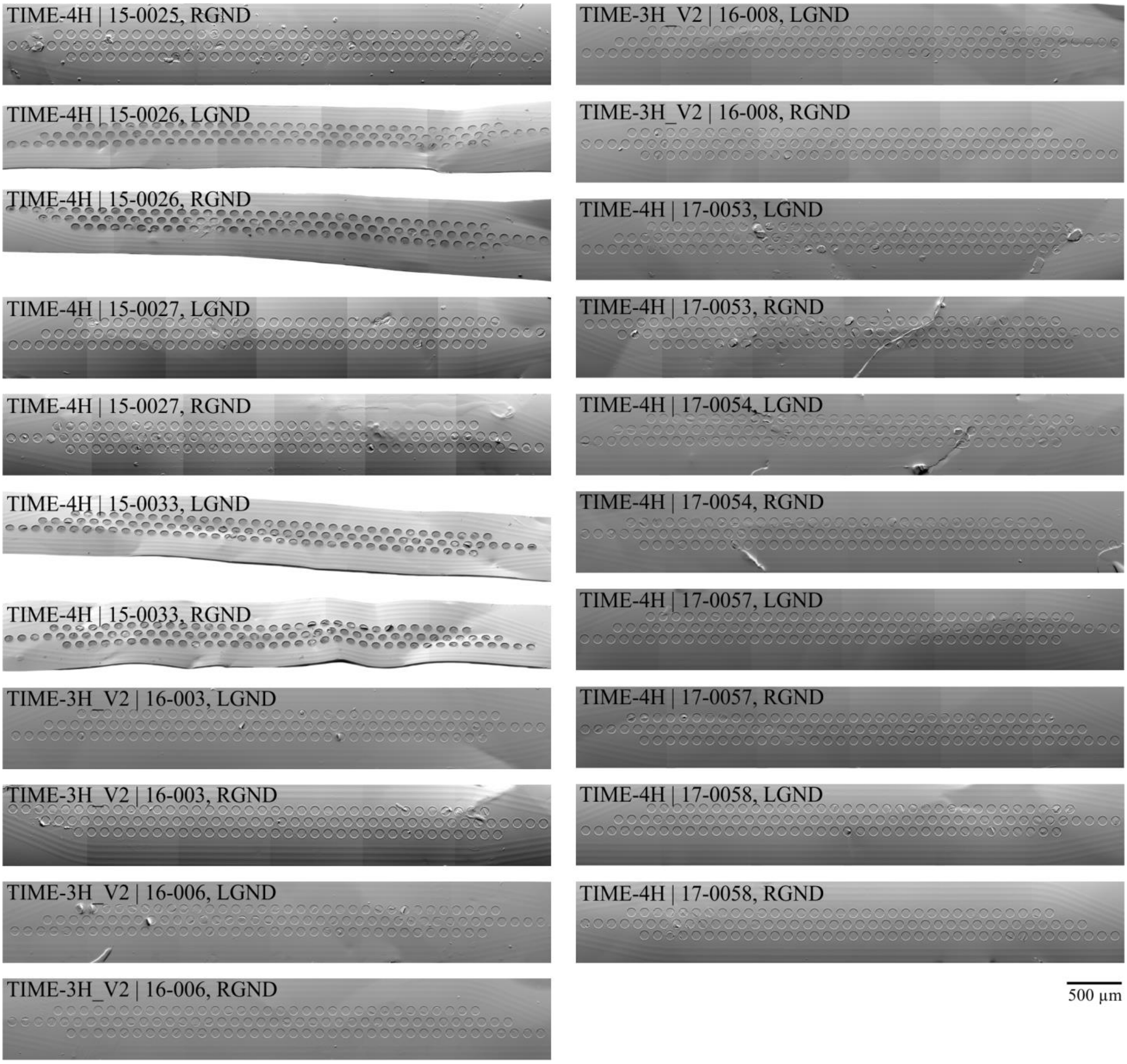


**Figure S2**: Overview of all explanted and for optical analysis available ground contact sites. Images were acquired with SEM.

[6] T. Guiho, D. Andreu, V.M. López-Alvarez, P. Čvančara, A. Hiairrassary, G. Granata, L. Wauters, W. Jensen, J.-L. Divoux, S. Micera, T. Stieglitz, X. Navarro, D. Guiraud, Advanced 56 Channels Stimulation System to Drive Intrafascicular Electrodes, in: J. Ibáñez, J. González-Vargas, J.M. Azorín, M. Akay, J.L. Pons (Eds.), Converging Clinical

and Engineering Research on Neurorehabilitation II, Springer International Publishing, Cham, 2017, pp. 743–747.

[7] L. Bisoni, C. Carboni, L. Raffo, N. Carta, M. Barbaro, An HV-CMOS Integrated Circuit for Neural Stimulation in Prosthetic Applications, IEEE Trans. Circuits Syst. II 62 (2015) 184–188. https://doi.org/10.1109/TCSII.2014.2387679.

[8] G. Valle, F.M. Petrini, I. Strauss, F. Iberite, E. D'Anna, G. Granata, M. Controzzi, C. Cipriani, T. Stieglitz, P.M. Rossini, A. Mazzoni, S. Raspopovic, S. Micera, Comparison of linear frequency and amplitude modulation for intraneural sensory feedback in bidirectional hand prostheses, Sci. Rep. 8 (2018) 16666. https://doi.org/10.1038/s41598-018-34910-w.

[9] S. Raspopovic, M. Capogrosso, F.M. Petrini, M. Bonizzato, J. Rigosa, G. Di Pino, J. Carpaneto, M. Controzzi, T. Boretius, E. Fernandez, G. Granata, C.M. Oddo, L. Citi, A.L. Ciancio, C. Cipriani, M.C. Carrozza, W. Jensen, E. Guglielmelli, T. Stieglitz, P.M. Rossini, S. Micera, Restoring natural sensory feedback in real-time bidirectional hand prostheses, Sci Transl Med 6 (2014) 222ra19. https://doi.org/10.1126/scitranslmed.3006820.

[10] G. Valle, A. Mazzoni, F. Iberite, E. D'Anna, I. Strauss, G. Granata, M. Controzzi, F. Clemente, G. Rognini, C. Cipriani, T. Stieglitz, F.M. Petrini, P.M. Rossini, S. Micera, Biomimetic Intraneural Sensory Feedback Enhances Sensation Naturalness, Tactile Sensitivity, and Manual Dexterity in a Bidirectional Prosthesis, Neuron 100 (2018) 37-45.e7. https://doi.org/10.1016/j.neuron.2018.08.033.
